## Supplementary data for "Characterization and deorphanization of RYamide signaling in *Aedes aegypti*: a potential regulator of hindgut-associated physiology"

**Supplementary Table S1. Information of designed primers.** Geneious Pro Bioinformatics Software was used to design primers.

| Primers | Sequences | Functions |
|---|---|---|
| AAEL017005-Kozak | TTCTGCCGCCACCATGAGCG | ORF cloning for functional receptor assay |
| AAEL017005-Stop | GGTTGTTGTATCACCGTAGC | |
| AAEL019786-Kozak | AAGCTTGCCACCATGAACTTCACTGCCGAG | ORF cloning for functional receptor assay |
| AAEL019786-Stop | ATTCTAGAATTCAACCCCCTACAACC | |
| RYa-qPCR-F | CTAATCCTTCTAGTCAGTGCGG | RT-qPCR amplification of *Aedae*RYamides |
| RYa-qPCR-R | AGCGGGATCCAAGAAAGAAGC | |
| RYa_RNAi-F | TGCGCCACCACACTCCATAT | dsRNA synthesis |
| RYa_RNAi-R | GGAGATCTTTCCCGCCGTAGA | |

**Supplementary Table S2. Primary amino acid sequences of *Ae. aegypti* neuropeptides used in immunohistochemistry and functional bioluminescence assay.** Highlighted sequences indicate the conservation of residues at the C-terminus.

| Name | Amino acid sequence |
|---|---|
| NPF | SFTDARPQDDPTSVAEAIRLLQELETKHAQHARPRFa |
| 2sNPF-1 | KAVRSPSLRLRFa |
| sNPF-1 | SPSLRLRFa |
| sNPF-2 | APQLRLRFa |
| sNPF-3 | APSQRLRWa |
| FMRFa-1 | SALDKNFMRFa |
| FMRFa-2 | ASKQANLMRFa |
| FMRFa-3 | AGQGFMRFa |
| FMRFa-4 | DSPKNLMRFa |
| FMRFa-8 | GSGNLMRFa |
| FMRFa-9 | AKGNLMRFa |
| FMRFa-11 | MDNNFMRFa |
| Head peptide 1 | (pE)-RP-(hP)-SLKTRFa |
| Myosuppressin | TDVDHVFLRFa |
| Pyrokinin1 (PK1) | AGNSGANSGMWFGPRLa |
| CAPA | pQGLVPFPRVa |
| RYamide 1 | PFFVGSRYa |
| RYamide 2 | NDRFFLGSRYa |

*Note: pE is pyro-glutamic acid; hP is 4-hydroxyproline*

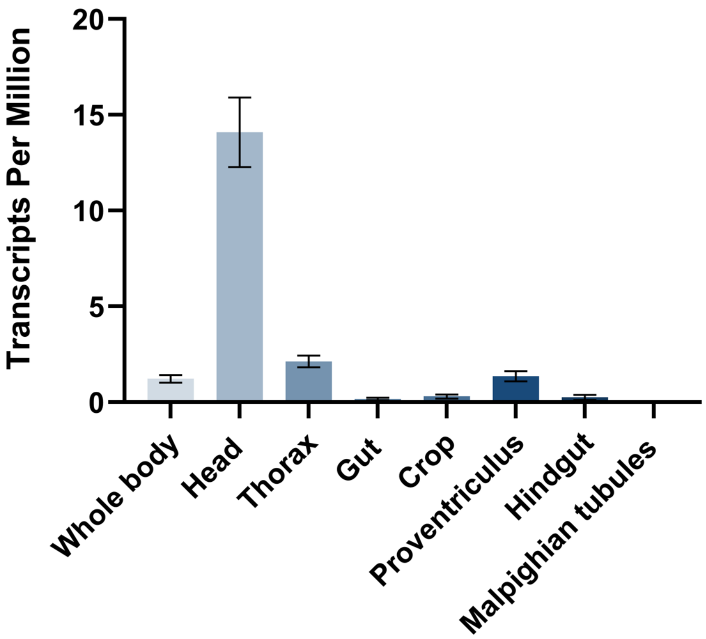


**Supplementary Figure S1. Organ-specific expression profile of RYamide in adult *Ae. aegypti* based on public RNAseq data.** Figure was prepared using data from Hixson *et al.* 2022[28].

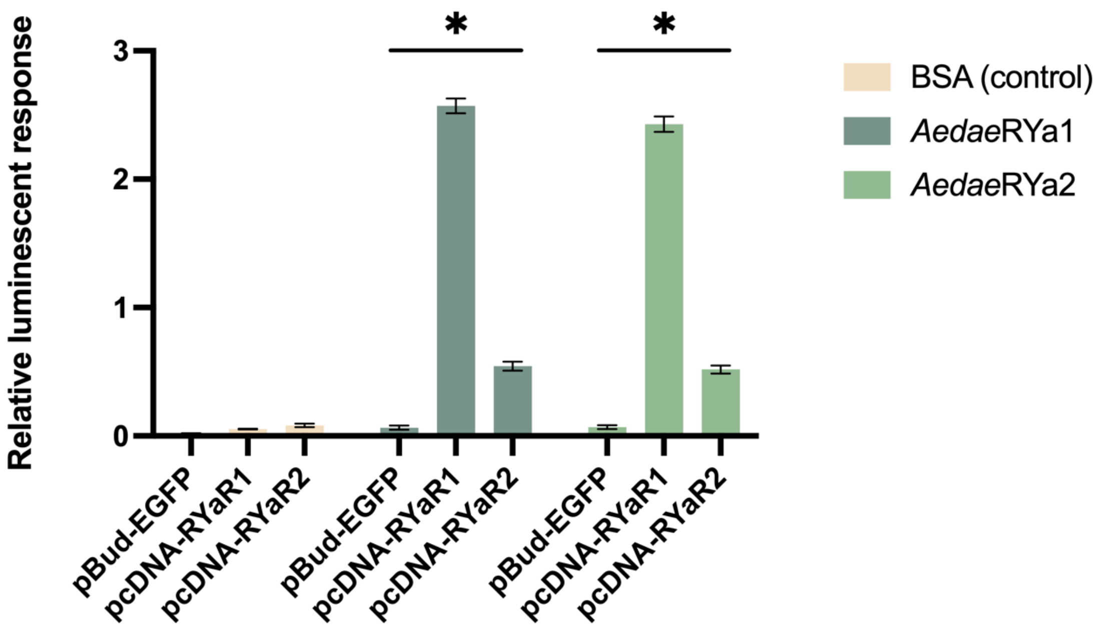


**Supplementary Figure S2. Normalized luminescent response of CHO-K1 cells expressing EGFP in pBud vector, *Ae. aegypti* RYaR2 (AAEL019786) in pcDNA3.1+ vector (n=3).** Assay media (BSA) alone and different ligands ($10^{-6}$M) were applied to cells expressing different expression constructs to validate the receptor activity by comparing the luminescent responses generated via receptor activation. Statistical differences are denoted with asterisk (*), as determined by a two-way ANOVA with Šídák's multiple comparisons test ($p<0.05$).

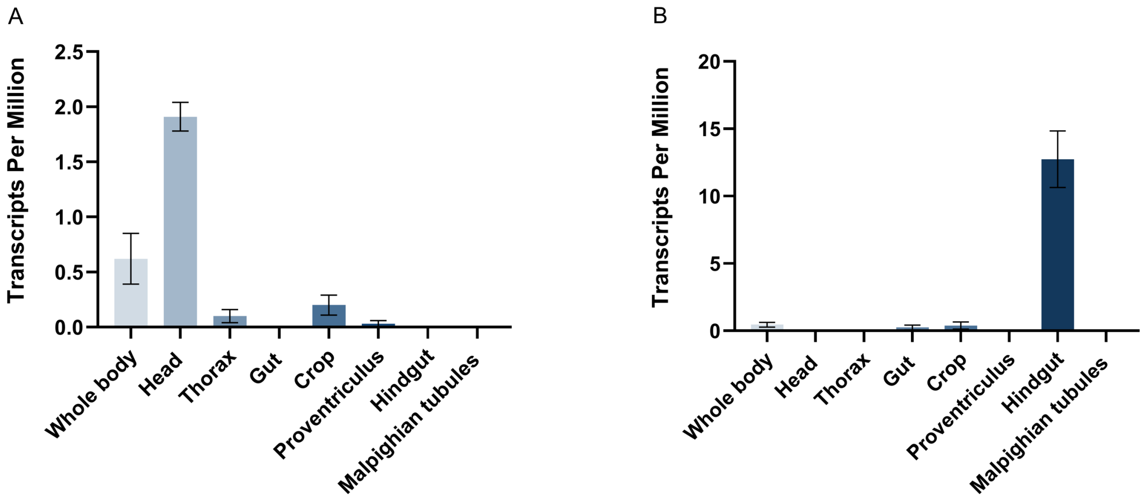


**Supplementary Figure S3. Organ-specific expression profile of RYaR1 (A) and RYaR2 (B) in adult *Ae. aegypti* based on public RNAseq data.** Figure was prepared using data from Hixson *et al.* 2022[28].

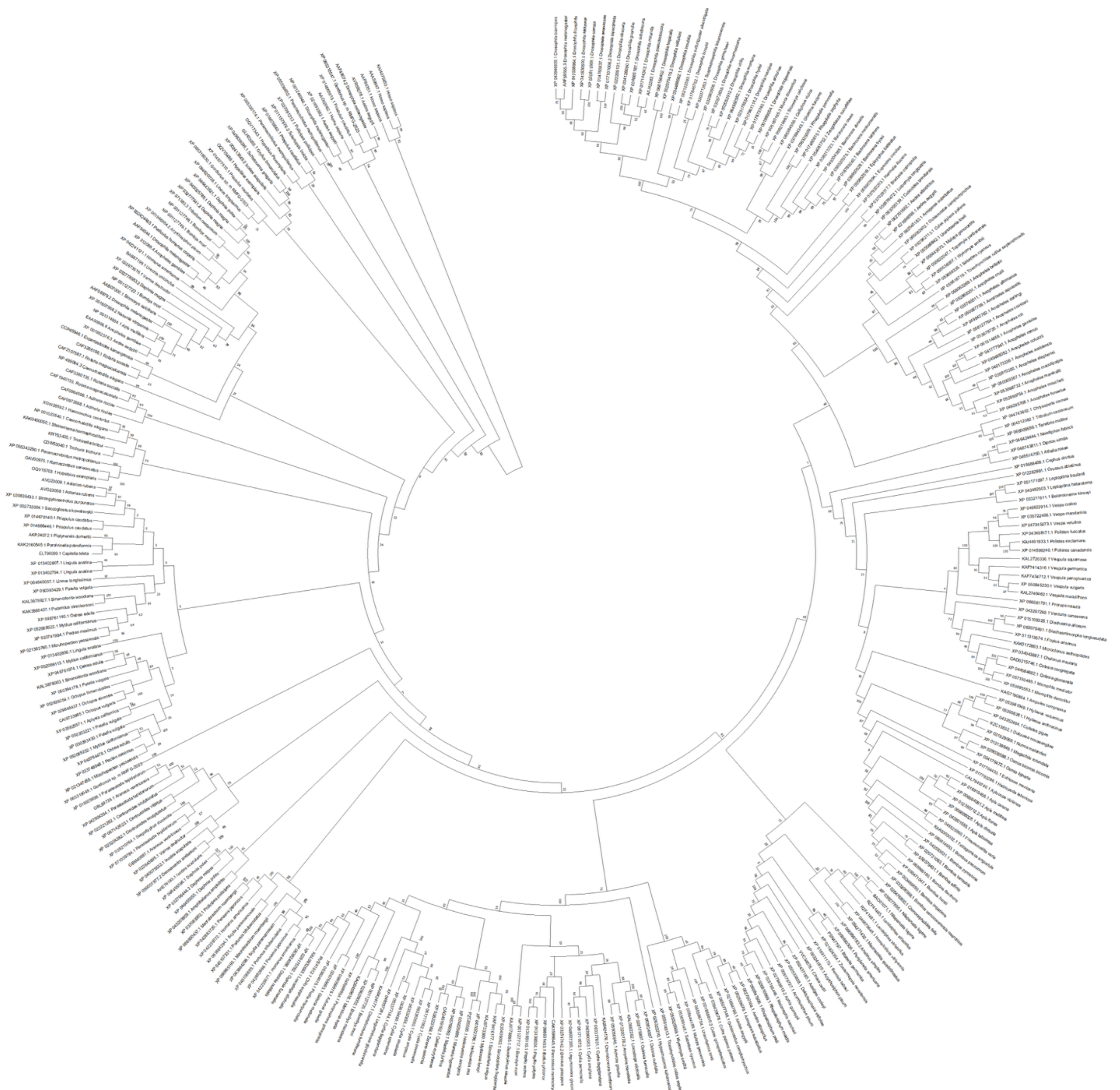

**Supplementary Figure S4. Phylogenetic relationship of RYamide receptors across protostomes and deuterostomes.** Tree was constructed using maximum-likelihood phylogenetic analysis methods (with 1000 bootstrap replicates). The annotated numbers adjacent to nodes indicate the support percentage for the clustering of related sequences within the respective clade. Tachykinin, NPF and sNPF receptors, as well as NPY receptors of *Homo sapiens*, were included to demonstrate the evolutionary relationship between vertebrate NPY receptors and invertebrate RYamide, tachykinin, NPF, and sNPF receptors. *Homo sapiens* NPYR was included in the analysis and imposed as the outgroup.
